# A Distinct Autofluorescence Distribution Pattern Marks Enzymatic Deconstruction of Plant Cell Wall

**DOI:** 10.1101/2024.12.12.628095

**Authors:** Solmaz Hossein Khani, Khadidja Ould Amer, Noah Remy, Berangère Lebas, Anouck Habrant, Ali Faraj, Grégoire Malandain, Gabriel Paës, Yassin Refahi

## Abstract

Achieving an economically viable transformation of plant cell walls into bioproducts requires a comprehensive understanding of enzymatic deconstruction. Microscale quantitative analysis offers a relevant approach to enhance our understanding of cell wall hydrolysis, but becomes challenging under high deconstruction conditions. This study comprehensively addresses the challenges of quantifying the impact of extensive enzymatic deconstruction on plant cell wall at microscale. Investigation of highly deconstructed spruce wood provided spatial profiles of cell walls during hydrolysis with a remarkable precision. A distinct cell wall autofluorescence distribution pattern marking enzymatic hydrolysis along with an asynchronous impact of hydrolysis on cell wall structure, with cell wall volume reduction preceding cell wall accessible surface area decrease, were revealed. This study provides novel insights into enzymatic deconstruction of cell wall at under-investigated cell scale, and a robust computational pipeline applicable to diverse biomass species and pretreatment types for assessing hydrolysis impact and efficiency.

## 1 Introduction

Plant cell wall, as a sustainable source of biopolymers, offers a renewable and carbon-neutral alternative to fossil-based resources [4, 41, 43, 46]. The shift towards utilizing plant cell wall biopolymers has the potential to significantly mitigate climate change, reduce our dependence on fossil carbon, and thus be considered as a cornerstone of a thriving bioeconomy [10, 36]. Plant cell wall is mainly composed of cellulose, hemicelluloses, lignin, and pectins, collectively referred to as lignocellulose [39]. Biotechnological processes for lignocellulose conversion involve enzymatic hydrolysis as a crucial and costly step, which is also known as saccharification [24]. During enzymatic hydrolysis, the enzymes break down the plant cell wall polysaccharides into fermentable sugars (primarily glucose) [21]. The critical challenge in sustainable transformation of plant cell wall is overcoming the inherent resistance of plant cell wall to deconstruction, called recalcitrance [19]. The recalcitrance contributes substantially to the high cost of bioproducts obtained from plant cell wall. Nanoscale recalcitrance markers, such as cellulose crystallinity, cell wall porosity, and lignin content [28, 45, 47, 48], have been the primary focus of research in recent decades. Immunolabeling, cellulose staining, and fluorescence imaging together with fluorescent probes have also been used to observe cell wall polymer deconstruction and characterize enzymatic hydrolysis dynamics through time-lapse imaging, providing visual evidence of cellular changes during enzymatic deconstruction [25, 44]. Despite these advances, morphological characteristics underlying the enzymatic deconstruction of plant cell wall at microscale, more precisely at cell and tissue scales, remains quantitatively under-investigated. By contrast, quantitative studies of cell growth and division, which can be considered the reverse of deconstruction, have yielded valuable insights by combining 4D (space + time) imaging and image processing [1, 14, 22, 40, 42]. The knowledge gap in understanding cell and tissue scales deconstruction of plant cell wall is mainly due to challenges in acquiring quantitative data on cell wall deconstruction dynamics at these scales. Moreover, the image processing methods employed in cell growth and division studies [15, 17, 20, 31, 37] are not suited to address the specific challenges of investigating plant cell wall deconstruction at cell and tissue scales. While the natural autofluorescence of cell wall facilitates the use of fluorescence confocal microscopy for capturing time-lapse images of cell wall enzymatic deconstruction [13], this process is technically demanding. It requires maintaining a constant optimal temperature (typically 50°C) for enzymatic reactions while also preventing evaporation of the enzymatic cocktail [50]. Once acquired, quantitative analysis of collected time-lapse 3D images is challenging due to specific data properties: i) sample is often not aligned with the 3D acquisition axes resulting in a tilted imaged sample; ii) sample moves relative to the microscope objective over imaging period; iii) sample material loss occurs as a result of enzymatic deconstruction sometimes manifesting as complete disintegration of the cell walls; iv) deconstruction, high temperature, and changes in cell wall mechanical properties induce sample deformation over time; v) cell wall autofluorescence intensity decreases over time due to the changes in the lignin environment during enzymatic hydrolysis and photobleaching due to fluorescence imaging [50].

To study cell wall enzymatic deconstruction at microscale, a method has recently been developed to track cell wall deconstruction from time-lapse 3D images of cell wall enzymatic hydrolysis [38]. This method involves initially segmenting the image of the sample prior to deconstruction, at a stage when the cell walls are intact. Using temporal propagation of spatial information [2], this pre-hydrolysis image segmentation is then propagated to generate segmentations of images acquired during deconstruction. Using this method, it was demonstrated that cell and tissue scale parameters can predict the hydrolysis yield, thus can serve as indicators of the efficiency of cellulose conversion to glucose. This study demonstrated the relevance and strengths of studying enzymatic deconstruction at cell and tissue scales and introduced a new method to track deconstruction at these scales. However, this method requires that the time-lapse images meet some prerequisites: the imaged sample needs to be flat and untilted so that deviations from these conditions result in under-segmentation errors. The acquisition of such “perfect” datasets is particularly challenging and labor-intensive, as a substantial number of collected datasets with misaligned samples have to be discarded. Additionally, since the segmentation of images of the deconstructed sample relies on propagating the pre-hydrolysis image segmentation using image registration, the quality of this registration, therefore segmentations, is compromised by substantial deconstruction and deformations, which limit the application of the method. This limitation is particularly critical in the context of the saccharification process whose objective is to achieve high efficiency in converting plant biomass through extensive deconstruction. Besides the natural imperfections of sample and deconstruction extent, the sample displacement, also known as sample drift, needs to be addressed. Sample drift is a prevalent challenge in nanoscale imaging, as single-molecule localization microscopy, which requires to be estimated and compensated [23, 26]. A common approach to drift compensation involves using fiducial markers, such as gold nanoparticles or fluorescent beads to track their positions and measure three-dimensional drift [5, 7, 11]. Marker-free drift correction approaches are also developed with the advantage of requiring no special sample preparation or modifications to imaging system [3, 12, 16, 26]. While in nanoscale imaging, the drift needs to be corrected to ensure high spatial resolution, in microscale imaging of cell wall deconstruction, the drift during individual acquisitions is negligible, however the sample displacement between acquisitions needs to be taken into account. This sample displacement over time-lapse imaging time (here 24 hours), together with material loss, cell walls deformations and autofluorescence intensity reduction requires to go beyond simple drift correction (which would involve a rigid transformation) to achieve reliable tracking. Essentially, the previously mentioned prerequisites are hard to fulfill, and under high deconstruction conditions, virtually impossible to achieve.

In this study, a robust marker-free pipeline, named HydroTrack (Hydrolysis robust Tracking), is proposed to quantify cell wall enzymatic hydrolysis under high deconstruction conditions at cell and tissue scales from the time-lapse 3D images without requiring any prerequisite. HydroTrack effectively handles tilted acquisition of non-flat samples subject to moving, and undergoing substantial deconstruction. HydroTrack was applied to time-lapse 3D image datasets of enzymatic deconstruction of pretreated spruce wood samples. The quantifications are then used to investigate cell wall deconstruction at microscale and study the impact of enzymatic hydrolysis on cell wall morphological parameters and cell wall autofluorescence. Spruce was selected due to the significant impact of climate change on European forests, particularly the notable instances of spruce dieback [33, 34] highlighting the need to valorize this biomass feedstock.

## 2 Materials and methods

### 2.1 4D imaging of cell wall enzymatic deconstruction via confocal microscopy

Spruce (*Picea abies*) wood was provided by INRAE, Grand-Est - Nancy, Champenoux, France. Fragments were cut from 2 cm side wood blocks into pieces measuring 0.5 cm in width, 0.5 cm in thickness, and 1 cm in length using razor blades. Sodium chlorite pretreatment was applied to the fragments using 1.25 g NaClO_2_ and 150 *µ*L acetic acid in 40 mL water at 70°C for one hour, repeated several times until delignification was achieved. Lignin content was determined using acetyl bromide assay procedure [32]. Pretreatment is required to improve hydrolysis of cell wall carbohydrates [6, 27] and sodium chlorite pretreatment is an effective method in lignin removal which increases digestibility of cell wall carbohydrates [18]. After drying, pretreated samples were sectioned into 40 *µ*m transverse slices using a microtome (Thermo Scientific, USA) with disposable blades. Sections were mounted in 60 *µ*L of enzymatic solution in acetate buffer (50 mM, pH 5) between cover glass and a gene frame (65 *µ*L, Thermo Scientific, Waltham, USA). Control time-lapses were acquired using buffer solution without enzymes. The microscopy slide was put inside the incubation chamber (microscope adapted, OKOlab, Italy), heated at 50°C, as the optimal temperature for the action of enzymes, and placed on the microscope stage [50]. For enzymatic solution, an enzymatic cocktail from IFPEN (Rueil-Malmaison, France) was used. This cocktail, derived from *Trichoderma reesei* RUT-C30, a strain of cellulolytic filamentous fungi, contained mainly cellulases and xylanases. The cocktail was diluted in acetate buffer to reach a final cellulase activity of 30 FPU/g of biomass. To capture the dynamics of plant cell wall deconstruction, using cell wall natural autofluorescence [13], time-lapse imaging was performed using a confocal laser scanning microscope. A 3D image (z-stack) was acquired every hour during 24 hours, using a 20× objective with a 2× zoom, a 0.8 *µ*m numerical aperture, and a scanning speed of 4 *µ*s/pixel on the Fluoview FV3000 confocal microscope (Olympus, Tokyo, Japan), generating a time-lapse of twenty five 3D images. The excitation laser was set to 405 nm with a detection range from 415 to 515 nm. The acquisitions (z-stacks) included 400 2D slices, each with a resolution of 512 × 512 pixels and a z-step of 0.3 *µ*m. Hereafter, the initial time-lapse image, capturing the cell walls while still intact, is referred to as the pre-hydrolysis image. Subsequent images are referred to as hydrolysis phase images. A time-lapse dataset collected with the enzymatic cocktail is referred to as a hydrolysis dataset in contrast to the control dataset as a time-lapse dataset collected without enzymes. The hydrolysis and control datasets were each collected in triplicate, resulting in three control time-lapse 3D images and three hydrolysis time-lapse 3D images.

### 2.2 4D image processing

Quantification of dynamics of cell wall autofluorescence intensity and morphological parameters at cell and tissue scales from time-lapse image datasets, requires tracking and segmentation. These parameters can serve as predictors of cell wall enzymatic hydrolysis efficiency [38]. To quantify the dynamics of these parameters, the challenges of processing time-lapse 3D images of highly deconstructed samples due to sample misalignment, drift, deformation, material loss, and reducing autofluorescence intensity over time, need to be addressed, particularly as these challenges intensify in extensive deconstruction conditions. To address these challenges, a tracking and segmentation pipeline named HydroTrack is developed. HydroTrack uses a combination of divide-and-conquer strategy [9] and temporal propagation of spatial information strategy [2, 38] to track cell wall deconstruction and identify individual cell walls from a time-lapse 3D image dataset. This is achieved by dividing the time-lapse images into sequential clusters and limiting image registration (needed for sample displacement and deformation compensation) to images inside each cluster. The resulting transformations are then combined to track the changes occurring over large temporal intervals. The transformations are also used to propagate the pre-hydrolysis cell walls along the hydrolysis phase time-lapse images. These time-constrained transformations are also used to identify individual cells in hydrolysis phase images using their initial state in pre-hydrolysis image via a propagation strategy.

#### 2.2.1 Sample displacement and deformation compensation

Sample displacement in x, y, or z directions between consecutive acquisitions, together with cell walls deformations and material loss during enzymatic deconstruction, can introduce inconsistencies in the imaged region. Consequently, the imaged region may vary between acquisitions within the same time-lapse dataset, complicating precise intensity dynamics analysis. Therefore, identifying the consistently imaged region across all time-lapse acquisitions as the part of the sample that stays within the microscope’s field of view regardless of sample displacement and alterations, is crucial for ensuring that the quantifications are accurate and comparable over time. Identification of the consistently imaged region requires registration of the images of the time-lapse dataset to compensate for deformation and displacement. To achieve this, two strategies can be employed. Let 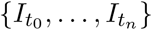 denote the dataset of time-lapse 3D images of cell wall deconstruction, where 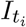 represents the *i*-th 3D image (z-stack) acquired at time *t*_*i*_, 0 ≤ *i* ≤ *n* and 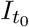 corresponds to the pre-hydrolysis image. Using 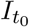, as the reference image for displacement and deformation compensation, two existing strategies can be employed: i) the *global* strategy which involves co-registering 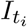, 0 *< i* ≤ *n* with the reference image, 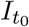, thus computing directly the transformation to resample 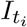 in the frame of 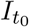; ii) the *local* strategy which involves co-registering each pair of successive images 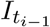 and 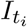, 0 *< i* ≤ *n* and composing the resulting transformations to obtain the cumulative transformation required to resample 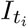 in the frame of 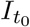. On the one hand, extensive deconstruction which implies significant changes between images, compromises the registration between temporally distant images, as it fails to accurately track substantial alterations generated by extensive enzymatic deconstruction. Additionally, intensity-based image registration may misinterpret shape changes with intensity variations, particularly if the intensity changes are substantial. Consequently, images that are temporally too distant should not be registered directly. On the other hand, composing transformations results in accumulation of registration errors and inaccuracies.

To address these challenges, HydroTrack employs a divide-and-conquer strategy, specifically devised for high deconstruction conditions, to constrain image registration to images within the clusters while minimizing the number of required transformations. This strategy involves dividing time-lapse images into sequential clusters, where each pair of successive clusters overlaps by one image: the last image of one cluster is also the first image of the next (Figure 1.A). More precisely, HydroTrack divides the sequence of 3D images into sequential clusters *C*_*q*_ 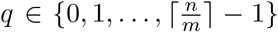. Each cluster *C*_*q*_, consists of *m* + 1 consecutive images acquired at {*t*_*qm*_,…,*t*_min(*q*+1)*m*,*n*)_}.i.e. 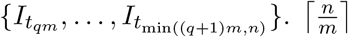 denotes the ceiling of 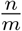, which is the smallest integer greater than or equal to 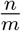. If *n* is not an exact multiple of *m*, the final cluster is formed with fewer than *m* + 1 images.

**Fig. 1:**
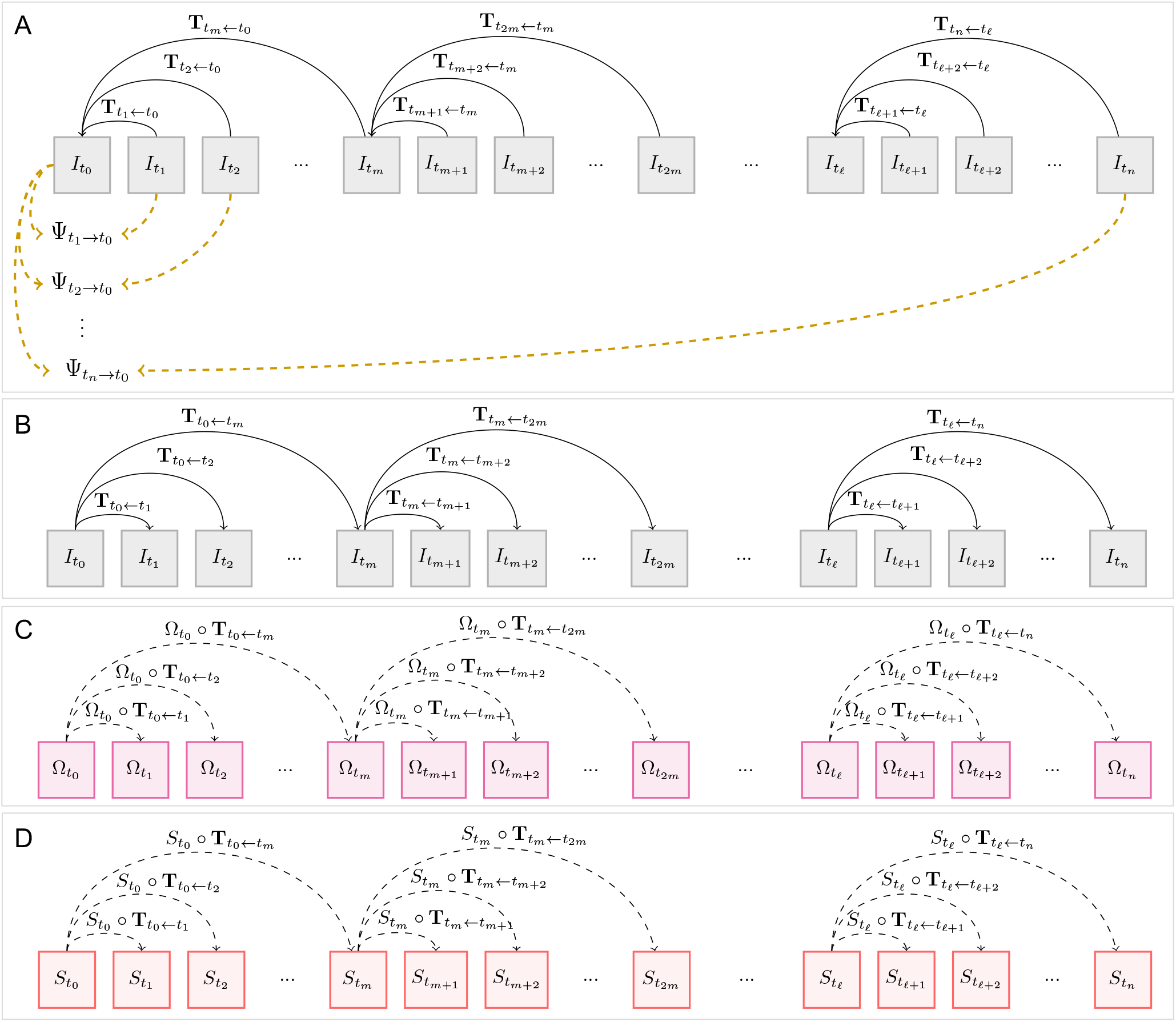
Tracking cell wall autofluorescence intensity during enzymatic hydrolysis and segmentation of hydrolysis phase images. 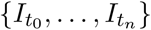 denotes the set of time-lapse 3D images, where 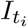 represents the *i*-th 3D image acquired at time *t*_*i*_, 0 ≤ *i* ≤ *n*. The first image, 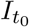, is called pre-hydrolysis image and the subsequent images are called hydrolysis phase images. (A) Identification of the consistently imaged region in the frame of pre-hydrolysis image; The time-lapse images are divided into sequential clusters of *m*+ 1 successive images with the first cluster starting from the pre-hydrolysis image. The last cluster includes the sequence of images 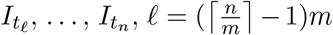. The intra-cluster images are registered with the first image of the cluster. The resulting transformations are then used to compute the tracked regions between pairs of pre-hydrolyis image, 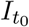, and the hydrolysis phase images, 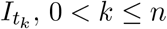 in the frame of the prehydrolysis image, denoted by 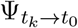. The consistently imaged region in the frame of the pre-hydrolysis image, denoted by 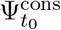, is the intersection of these tracked regions 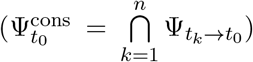. (B) The intra-cluster images are registered with the last image of the cluster. (C) Cell wall mask of consistently imaged region in the frame of pre-hydrolysis image, denoted by 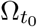, is propagated to subsequent time points using the resulting intra-cluster forward transformations to compute cell wall masks of consistently imaged regions in the frame of hydrolysis phase images, denoted by 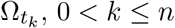. (D) The intra-cluster forward transformations are also used to propagate pre-hydrolysis cell segmentation, denoted by 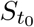, to subsequent time points to generate cell segmentation of hydrolysis phase images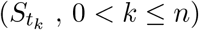.

After dividing the images into clusters, transformations are computed within these clusters in the backward direction, enabling the registration of images within each cluster with the initial image of that cluster (Figure 1.A). Hereafter, the transformation that registers the floating image, 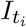, with the reference image, 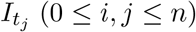, and resamples 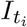 in the frame of 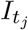, is denoted by 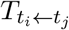. The resampled image is obtained through application of the computed transformation to the floating image 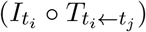 and is denoted as 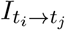. Resampling requires interpolation. For autofluorescence intensity images, linear interpolation is employed, whereas nearest-neighbor interpolation is used for label images (binary masks and segmentations). This registration process begins by first computing the rigid transformation to deal with translations and rotations between images. This rigid transformation is then used to compute an affine transformation which is, in turn, used to compute the non-linear transformation. All these registrations used the block matching framework [35].

The resulting backward transformations are then used to resample the hydrolysis phase images, 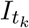 (for 1 ≤ *k* ≤ *n*), in the frame of pre-hydrolysis image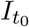. To do this, given a hydrolysis phase image, 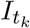, the left boundary image of the cluster in which 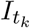 resides is identified 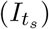. The transformation resulting from registering 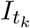 to 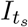 is used to resample 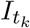. This initially resampled image is subsequently resampled by sequential application of the transformation resulting from registration of the boundary images of the preceding clusters. To be more precise, if *k* ≤ *m*, the resampled image, denoted as 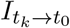, is given by 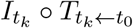. If 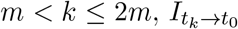 is computed as 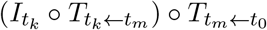. If 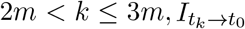, is computed by 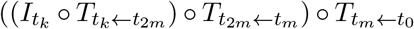. More generally, for 0 *< k* ≤ *n*, the resampled image 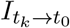 is computed by the following recursive application of transformations: 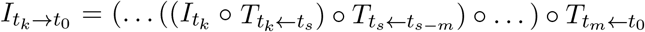, where 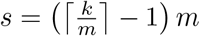

Following the resampling of the hydrolysis phase images in the frame of the pre-hydrolysis image, HydroTrack proceeds with the following steps: i) identifying the consistently imaged region in the frame of pre-hydrolysis image; ii) generating a cell wall mask of this region to eliminate background noise and propagating this mask to track the initial cell walls evolution throughout the imaging time. The Sections 2.2.2 and 2.2.3 provide detailed descriptions of these steps.

#### 2.2.2 Identification of consistently imaged region

Detecting the portion of the sample which stays within the objective field of view despite sample drift, referred to as the consistently imaged region, to restrict the quantification to this region, is essential for ensuring that the quantifications are accurate and comparable over time. To determine the consistently imaged region, HydroTrack first identifies the temporally consistent imaged regions between pairs of pre-hydrolysis and hydrolysis phase images. The resampling of hydrolysis phase images in the frame of the pre-hydrolysis image 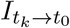 (for 1 ≤ *k* ≤ *n*), as described previously, enables the identification of these pair-wise temporally consistent regions. For each resampled image 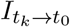, HydroTrack generates a binary mask 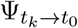. Each voxel (3D pixel) of this mask is set to 1 if it maps back to a voxel inside the corresponding hydrolysis phase image 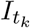, and 0 otherwise (Figure 1.A). In simpler terms, this binary mask indicates the part of the pre-hydrolysis image covered by the resampled hydrolysis phase image 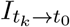. The consistently imaged region in the frame of the pre-hydrolysis image, denoted by 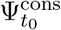, is then defined as the intersection of these binary masks and computed using a logical AND operation:

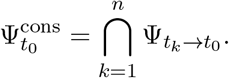

#### 2.2.3 Propagation-based cell wall tracking

Following identification of the consistently imaged region, it is essential to isolate the cell walls within this region of the pre-hydrolysis image to discard the background, which can be noisy and can interfere with accurate intensity measurements. This is achieved by generating a cell wall mask within 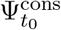 using thresholding of pre-hydrolysis image. The threshold value, set to 340 and determined by an expert, is used uniformly for all of the collected time-lapse datasets. In the resulting binary mask, denoted as 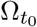, to each voxel one of the two distinct values is attributed. These values represent the background and the foreground corresponding to the cell walls of the pre-hydrolysis consistently imaged region. After computing 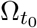, which indicates the pre-hydrolysis consistently imaged cell walls, it is essential to track this specific region over time to focus only on the originally identified cells walls 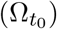. This is ensured by propagation of 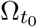 to identify and track the region marked as pre-hydrolysis cell walls throughout the entire time-lapse.

HydroTrack proceeds by using the divide-and-conquer registration strategy and computes the forward non-linear transformations required to register the starting image of each cluster with the subsequent images within that cluster (Figure 1.B). Similar to backward transformations, the block matching framework is used. Following these intra-cluster forward registrations, successive application of the transformations allows the propagation of 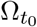. More precisely, let 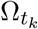 (0 *< k* ≤ *n*) denote the region mapping back to 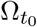, in the frame of 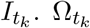 is obtained recursively as follows:

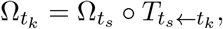

where 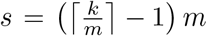, and 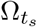 is the cell wall mask within the consistently imaged region of the starting image of the cluster number 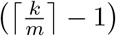 (Figure 1.C). 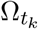 can be also expressed through successive applications of forward transformations starting from 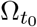

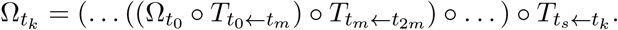

This expanded version involves transformations from registration of boundary images of all preceding clusters (crossing the clusters) ending with the transformation from a registration within the target cluster (which includes 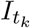).

This propagation-based cell wall tracking approach allows to focus on the initially identified cell walls and the corresponding regions at subsequent time points, thereby providing access to the dynamics of cell wall autofluorescence intensity. This maintains consistency and enables accurate and comparable assessment of cell walls deconstruction dynamics across the entire time-lapse dataset.

#### 2.2.4 Individual cell identification in time-lapse 3D images

Studying the cell wall enzymatic deconstruction at cell and tissue scales requires computing and analyzing morphological parameters dynamics during enzymatic deconstruction. These parameters include cell wall volume and cell wall accessible surface area which can be used as indicators of the cellulose conversion efficiency [38]. To achieve this, the time-lapse 3D images should be segmented and cells should be individually identified. HydroTrack achieves this by initially segmenting the pre-hydrolysis 3D image that may be misaligned with the imaging plane. The hydrolysis phase segmentation is performed using a propagation strategy. In this strategy, the forward intra-cluster transformations computed in Section 2.2.3 and shown in (Figure 1.B) are subsequently applied to propagate the pre-hydrolysis segmentation to compute hydrolysis phase image segmentations. The following provides a detailed description of the pre-hydrolysis image segmentation and the hydrolysis phase image segmentation.

### Pre-hydrolysis image segmentation

The spruce wood samples imaged during enzymatic deconstruction are cross-sectional slices of wood consisting of plant cells with open ends. To achieve a cell resolution segmentation of the pre-hydrolysis sample image, it is crucial to virtually close the open ends of these cells and isolate the sample from its exterior. This task is straightforward when the sample is flat and untilted, as virtual closures can be applied to the top and bottom of the sample, treating these boundaries as *xy* planes normal to the *z*-axis. However, for non-flat or tilted samples, effective use of the previously developed tools [38] requires trimming the upper and lower portions of the sample image to achieve a flat and untilted sample image. This adjustment, while essential for accurate segmentation, results in the exclusion of valuable portions of the sample image, particularly those parts that later undergo deconstruction. Therefore, limiting analysis to datasets of perfectly leveled acquisitions is essential but collecting such datasets is labor-intensive, since sample preparation and conditions of time-lapse imaging lead to many datasets of non-flat, and tilted samples that must be excluded, increasing the number of experiments needed to collect the desired leveled acquisitions.

HydroTrack successfully segments tilted sample acquisitions, therefore reduces significantly data collection efforts. To achieve this, HydroTrack first isolates the sample from its exterior in pre-hydrolysis image by calculating the 3D convex hull of the sample. To compute the 3D convex hull, a 3D binary mask of the sample is first computed by thresholding using an expert determined threshold value of 340, uniformly used for all datasets. Subsequently, the mask undergoes a series of refinement steps. First, the holes, background voxels enclosed by foreground voxels, are filled. A binary dilation algorithm is then applied to further refine the mask. Finally, the refinement is further extended by employing the Chan-Vese algorithm [8] which is a level-set like segmentation method. The 3D convex hull is determined based on the coordinates of the largest connected component area of the mask, removing any remaining noise. The vertices of the 3D convex hull are then extracted and connected using Delaunay triangulation, forming a boundary that encompasses the imaged sample. The *h*-minima operator, constrained within the convex hull, was then used to detect areas of local minimum intensity located within each individual cell. These labeled areas served as seeds for a subsequent watershed algorithm also constrained within the confocal acquisition region defined by the convex hull boundary. Without this spatial constraint, the watershed algorithm produces substantial under-segmentation errors caused by open-ended cells. To identify cell walls from the resulting cell-resolution segmentations, thresholding is applied with the threshold value set to 340. Alternatively, 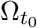 can be used to identify cell walls for each individual cell from the resulting cell-resolution segmentation discarding those out of the consistently imaged region. Figure 5.A illustrates an example of the computed 3D convex hull and the cell wall segmentation obtained by constraining the watershed algorithm to the region within the 3D convex hull, followed by thresholding. Hereafter, the segmentation obtained from pre-hydrolysis image through the watershed algorithm, denoted by 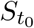, is referred to as ‘*cell segmentation*’, and the segmentation resulting from thresholding is referred to as ‘*cell wall segmentation*.’

### Hydrolysis phase image segmentation

After segmenting the pre-hydrolysis image, the hydrolysis phase segmentation is performed using a propagation strategy similar to that employed in the propagation-based cell wall tracking. In this strategy, the forward intra-cluster transformations computed in Section 2.2.3 and shown in (Figure 1.B) are sequentially applied to propagate the pre-hydrolysis segmentation generating segmentations of hydrolysis phase images (Figure 1.D). Let 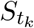 denote the cell segmentation at *t*_*k*_, 0 *< k* ≤ *n*. 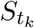 is obtained recursively as follows:

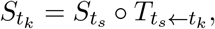

where 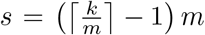, and 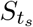 is the starting cell segmentation of the cluster number 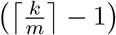. The expanded version expressing 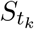 through successive applications of forward transformations starting from 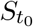 is the following:

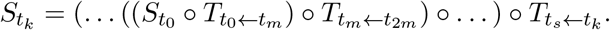

Similar to the cell wall propagation, this expanded version involves transformations from registering boundary images of preceding clusters (crossing the clusters) followed by a final transformation within the target cluste. Cell wall segmentations are then determined by thresholding with threshold value used for the pre-hydrolysis cell wall segmentation (expert determined value of 340) used uniformly for all images.

### Evaluation of segmentations and implementation

To evaluate the quality of segmentations generated by HydroTrack, Dice score is used which is a metric used for evaluating the accuracy and consistency of segmentation methods in image analysis. The Dice score, also known as the Sørensen–Dice similarity coefficient (DSC), is calculated as twice the overlap between the predicted segmentation (in our study segmentation obtained using HydroTrack) and the benchmark segmentation, divided by the total number of pixels in both the predicted and benchmark segmentations. More precisely, Dice score is defined as 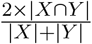 where *X* is the set of voxels in the segmentation obtained by HydroTrack, and *Y* is the set of voxels in the benchmark. The Dice score ranges from zero to one, with zero indicating no overlap and one indicating perfect overlap. After segmenting time-lapse images, HydroTrack computes cell wall volume by counting the number of voxels assigned the same label and multiplying this count by the voxel volume (0.116 *µ*m^3^ in this study). To determine the cell wall surface area, HydroTrack generates triangular meshes of the cell walls using the marching cubes algorithm. HydroTrack is implemented in Python 3 using Numpy, scikit-image, and SciPy, matplotlib, VTK packages.

## 3 Results and discussion

### 3.1 Robust quantification of cell wall enzymatic deconstruction at microscale

Studying the cell wall enzymatic deconstruction at microscale is a promising approach as cell and tissue scale parameters can be used to assess the efficiency of cellulose conversion into glucose [38]. To investigate spruce tree microscale enzymatic deconstruction, time-lapse datasets of pretreated spruce samples undergoing enzymatic hydrolysis together with control datasets were acquired. The hydrolysis datasets of possibly tilted samples exhibited highly deconstructed cell walls, especially at later time points (Figure 2). Furthermore, the hydrolysis datasets exhibited sample shifts relative to the microscope objective along with material loss, deformations and reducing cell wall autofluorescence intensity (Figure 4.A). To investigate the impact of enzymatic deconstruction on cell walls and compute microscale parameters dynamics from the collected time-lapse images, a robust 4D image processing pipeline, called HydroTrack, was developed (Figure 3). HydroTrack comprehensively overcomes various challenges of processing the time-lapse images due to sample misalignment, displacement, cell wall autofluorescence decrease, material loss, and deformation. It is specifically designed for high deconstruction conditions where these challenges intensify.

**Fig. 2:**
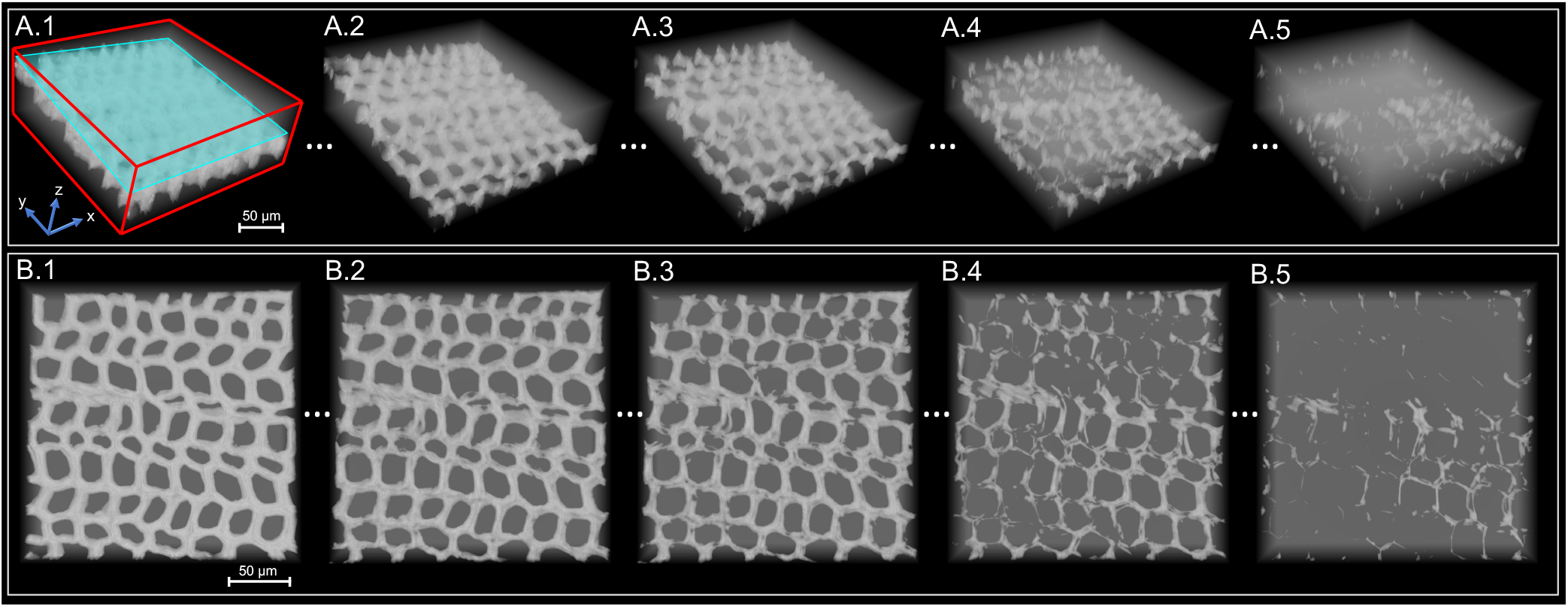
4D imaging of pretreated spruce wood sample during enzymatic hydrolysis captured using cell wall autofluorescence, with an enzymatic activity of 30 FPU/g of biomass. (A.1 – A.5) Side views of time-lapse 3D images shown at 6-hour intervals. The background is marked by a semi-transparent white color. In the first image, A.1, a cyan tangent plane on the sample surface and red lines are drawn to facilitate the visual distinction of the boundaries of the 3D images and to provide a visual reference for the spatial orientation of the wood sample in the 3D image. (B.1 – B.5) Top views of the time-lapse 3D images, shown at 6-hour intervals, with background marked by a semi-transparent white color. A.1 and B.1 show the pre-hydrolysis image and the subsequent images (A.2–A.5 and B.2–B.5) are hydrolysis phase images.

**Fig. 3:**
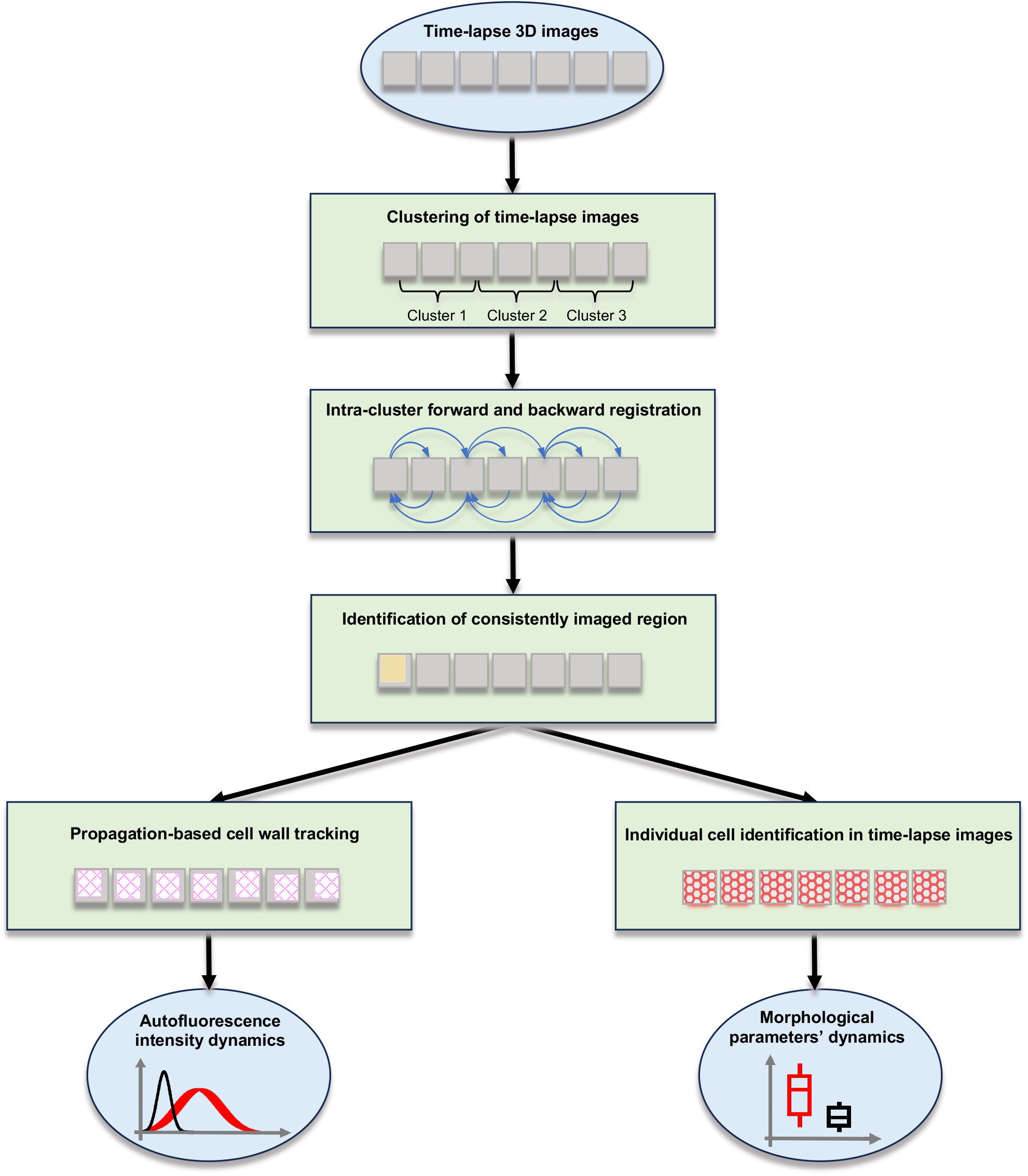
Schematic representation of the HydroTrack workflow. The workflow begins with input data which is a time-lapse 3D images of cell wall during enzymatic deconstruction. The time-lapse images is divided into clusters to improve the image registration quality. The intra-cluster images are registered bidirectionally: forward to the cluster’s last image and backward to its first image. The resulting transformations are used to track the cell wall autofluorescence intensity by first determining the consistently imaged region in pre-hydrolysis image and then by propagating pre-hydrolysis cell wall mask. The resulting transformations are also used to propagate the pre-hydrolysis image segmentation to generate the hydrolysis phase segmentations and identify individual cells in time-lapse images. Subsequently, HydroTrack provides autofluorescence intensity dynamics along with cell wall morphological parameters’ dynamics.

**Fig. 4:**
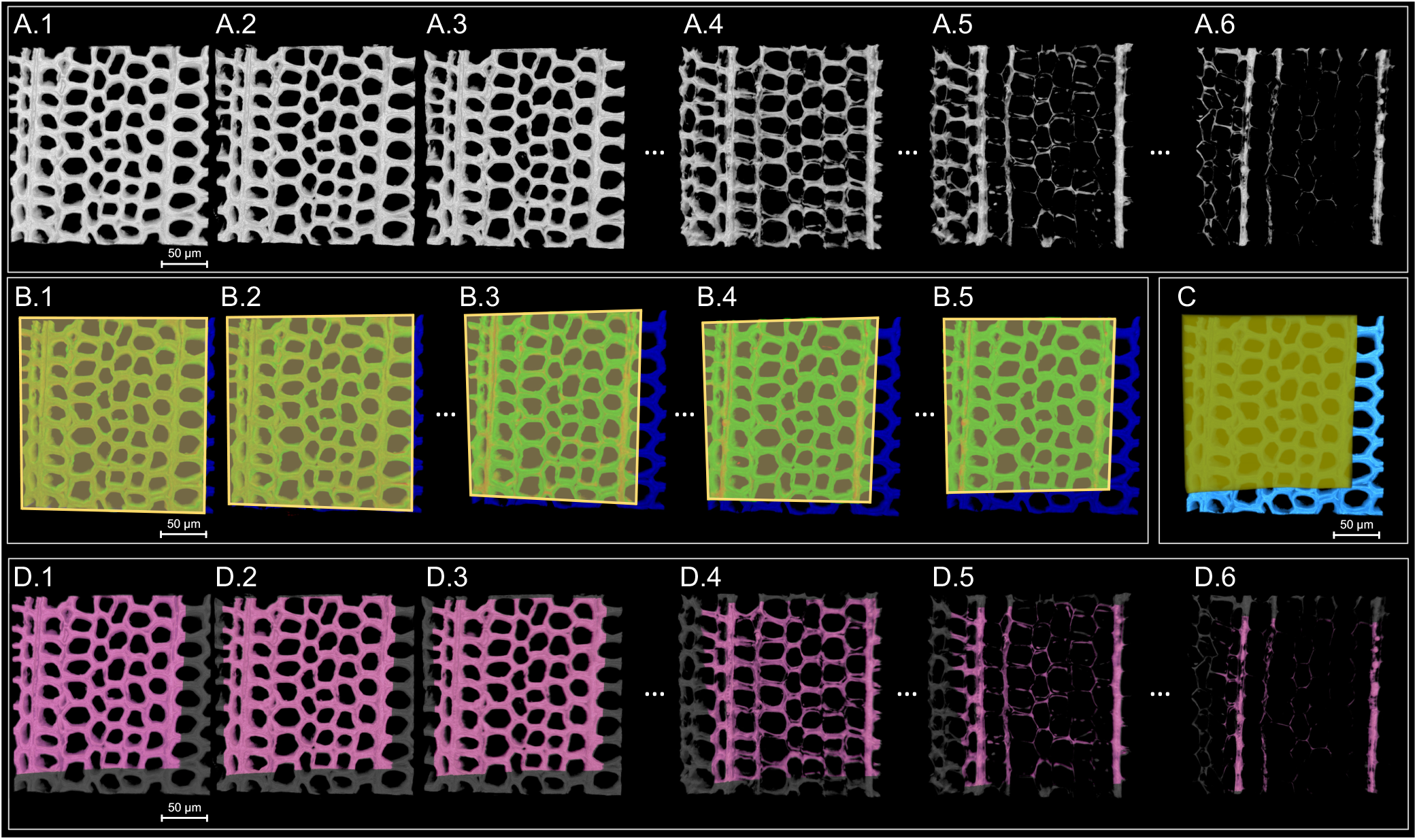
Cell wall tracking of pretreated spruce wood during enzymatic deconstruction. (A) Time-lapse 3D images of spruce wood enzymatic deconstruction with sample displacement between consecutive acquisitions. A.1 – A.6 correspond to 3D acquisitions at 0, 1, 2, 12, 18, and 24 hours. (B) Identification of tracked region between pairs of hydrolysis phase images (images at 1, 2, 12, 18, and 24 hours are shown) and pre-hydrolysis image in the frame of pre-hydrolysis image shown in A.1. These tracked regions are delimited by yellow quadrilaterals with a semi-transparent yellow fill. The quadrilaterals in B.1 – B.5 delimit the regions defined by 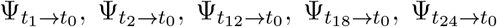, respectively, presented in Figure 1.A. The portion of the pre-hydrolysis image that was not captured in the hydrolysis phase image (therefore untracked) is marked in dark blue. (C) The consistently imaged region in the frame of the pre-hydrolysis image is marked by transparent yellow color. The remaining pre-hydrolysis image portion is marked in sky blue. (D) Cell wall masks of the consistently imaged regions are marked by magenta color and the remaining cell walls are marked in dark gray. D.1 – D.6 correspond to 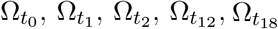, and 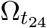 respectively, presented in Figure 1.C.

**Fig. 5:**
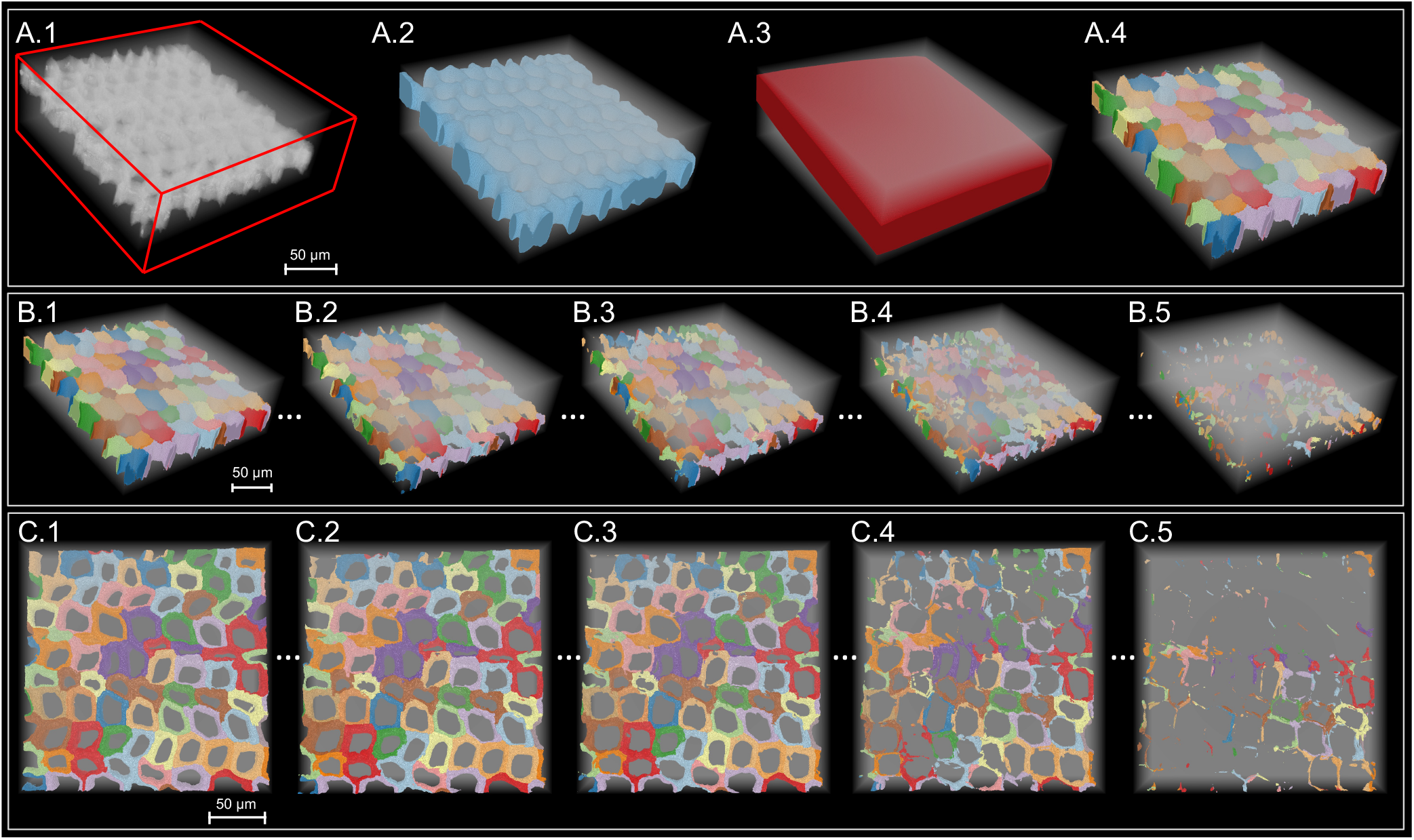
Segmentation of time-lapse images of pretreated spruce wood during enzymatic hydrolysis. (A.1 – A.4) Segmentation of pre-hydrolysis 3D acquisition of a tilted sample. (A.1) 3D image exhibiting a tilt. The red lines are included to enhance the visual clarity of the 3D image boundaries and to serve as a reference for the spatial orientation of the spruce sample within the 3D visualization. (A.2) 3D mask of the sample cell walls shown in blue. (A.3) 3D convex hull of the sample marked in red. (A.4) Successful application of the watershed algorithm to the region of the acquired 3D image constrained by the convex hull followed by thresholding allowing the identification of individual cell walls. In A – D, the semi-transparent white color marks the background. (B) Cell wall segmentations of time-lapse 3D images of a tilted sample, displayed at 6-hour intervals. The background appears in semi-transparent white, and cell walls are distinguished by lineage-specific coloring. (C) The same tilted sample’s cell wall segmentation and tracking, viewed from an overhead perspective. The cluster size is 7.

Temporal inconsistencies in the imaged sample region arise from displacement accompanied by cell wall deformations, cell wall autofluorescence intensity reduction and enzyme induced cell wall disintegration. This leads to variations in the imaged region across acquisitions in the same time-lapse dataset which requires an elaborate compensation beyond a conventional drift correction to identify consistently imaged region. The consistently imaged region is the portion of the sample that remains in the objective’s field of view over the course of imaging. The identification of consistently imaged region requires image registration either by registering hydrolysis phase images with a reference image as the pre-hydrolysis image (global strategy) or by sequentially registering pairs of successive images and composing the resulting transformations (local strategy). The global strategy fails under extensive deconstruction conditions, which introduce significant changes, particularly between distant images hindering registration. In addition, intensity-based registration can confuse shape changes with intensity variations when changes are substantial. Conversely, the local strategy suffers from progressive accumulation of registration errors and inaccuracies inherent in composing transformations. HydroTrack offers a compromise by dividing timelapse images into sequential clusters, and limiting the registration to intra-cluster images. Inter-cluster forward and backward registrations are then performed: the first image of each cluster is registered with other images within that cluster, and those images are, in turn, registered back to the first image. The registration of a hydrolysis phase image with pre-hydrolysis images involves successive application of transformations from backward registration of boundary images of preceding clusters of the hydrolysis phase image (crossing the clusters) ending with the transformation from a backward registration within the cluster including the hydrolysis phase image. The intersection of regions covered by all registered hydrolysis phase images with the pre-hydrolysis image (Figure 4.B) defines the consistently imaged region (Figure 4.C). This balanced solution limits the temporal distance between images for registration, while reducing the required number of transformations and can effectively reduce the cumulative error associated with the frequency of registrations and transformation applications typical of the local strategy while ensuring precise registration compared to global strategy. This can be also viewed as a hybrid approach combining the global strategy applied to intra-cluster images and a cluster-resolution local strategy involving the boundary images of clusters.

Following the identification of the consistently imaged region, HydroTrack ensures precise and consistent tracking of cell walls throughout the course of hydrolysis. This is achieved by first identifying the consistently imaged cell walls in the pre-hydrolysis image. A propagation strategy is then applied, using forward intra-cluster transformations to track these cell walls over the hydrolysis phase images (Figure 4.D). The cell wall propagation toward a given hydrolysis phase image involves a sequence of transformations: first, a series of transformations derived from registering boundary images of preceding clusters, and second, a transformation resulting from registration of the initial image of the target cluster (containing the hydrolysis phase image) with the hydrolysis phase image.

In addition to cell wall tracking which enables tracking cell wall autofluorescence intensities over the course of imaging, HydroTrack provides dynamics of cell wall morphological parameter which are indicators of cellulose conversion efficiency [38]. This is achieved by first segmenting the pre-hydrolysis which involves computing a convex hull of a potentially tilted sample within pre-hydrolysis image and applying the watershed algorithm within the boundaries defined by the convex hull (Figure 5.A). The cell wall disintegration, in particular under extensive deconstruction conditions, makes detecting individual cells in hydrolysis phase images challenging. In addition, enzymatic deconstruction induces a detachment between the compound middle lamella and the secondary cell wall [49]. To track the individual cells over the course of hydrolysis, a forward propagation strategy is applied which enables to identify the individual hydrolysis phase cells using their pre-hydrolysis state. This propagation is essentially similar to that used for tracking cell walls. A thresholding is then uniformly applied to separate individual cell walls from lumens (Figure 5.B & C).

To investigate the impact of cluster size and segmentation resampling frequency, the segmentations obtained with different cluster sizes were compared by computing Dice scores with benchmark segmentations obtained using a previously developed method [38] and corrected through visual inspection. The results indicated that the smaller the cluster size the lower the Dice score (Figure 6). Specifically, as the cluster size increased, Dice scores tended to concentrate around higher values, indicating improved similarity. The Dice score distribution for cluster size of 7 (*m* = 6) were notable, exhibiting a concentration of scores close to one, compared to smaller cluster sizes. These results highlight the importance of HydroTrack’s balanced approach offering a compromise by restricting temporal distance between two images to register and reducing the resampling frequency which enables to achieve accurate results.

**Fig. 6:**
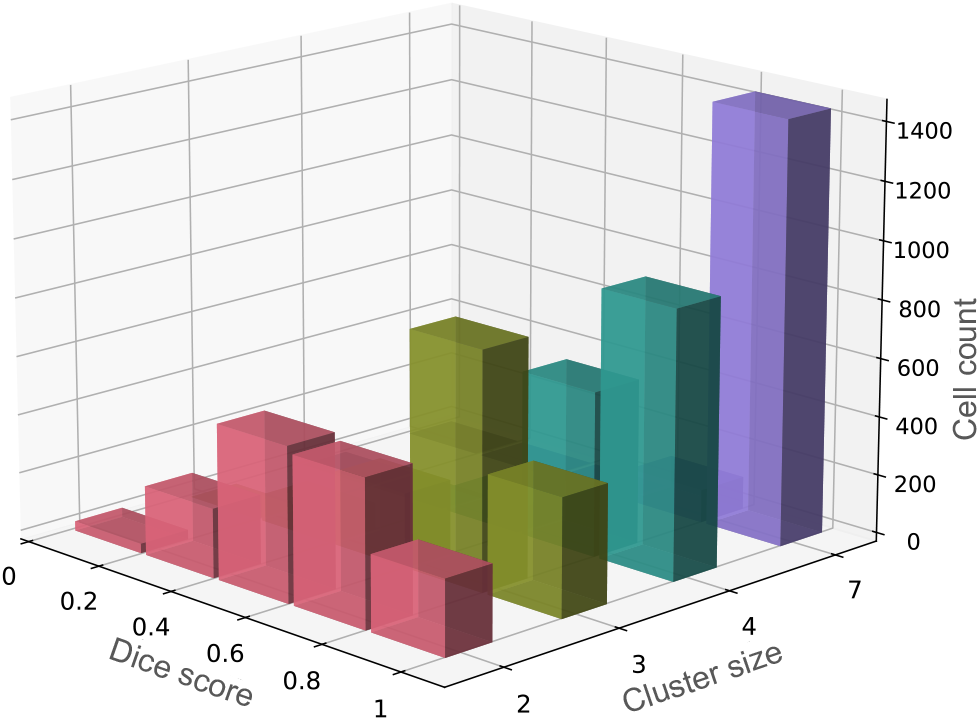
Distribution of Dice scores used to evaluate the effect of resampling frequency and segmentation quality between the segmentations generated by HydroTrack and the benchmark segmentations. Results are shown for cluster sizes of 2, 3, 4, 7. Dice score ranges from zero (for no overlap) to one (for perfect overlap).

### 3.2 Spatial profiling of extensive cell wall enzymatic deconstruction

Under high deconstruction conditions substantial changes occur between time-lapse images of plant cell wall enzymatic deconstruction. These significant changes introduced by enzymatic deconstruction and imaging conditions, challenge image registration accuracy and cause the registration of temporally distant images to fail (Figure 7.A.1, A.2, & A.3). The failure to precisely register temporally spaced images makes the accurate spatial analysis of deconstruction unreachable.

**Fig. 7:**
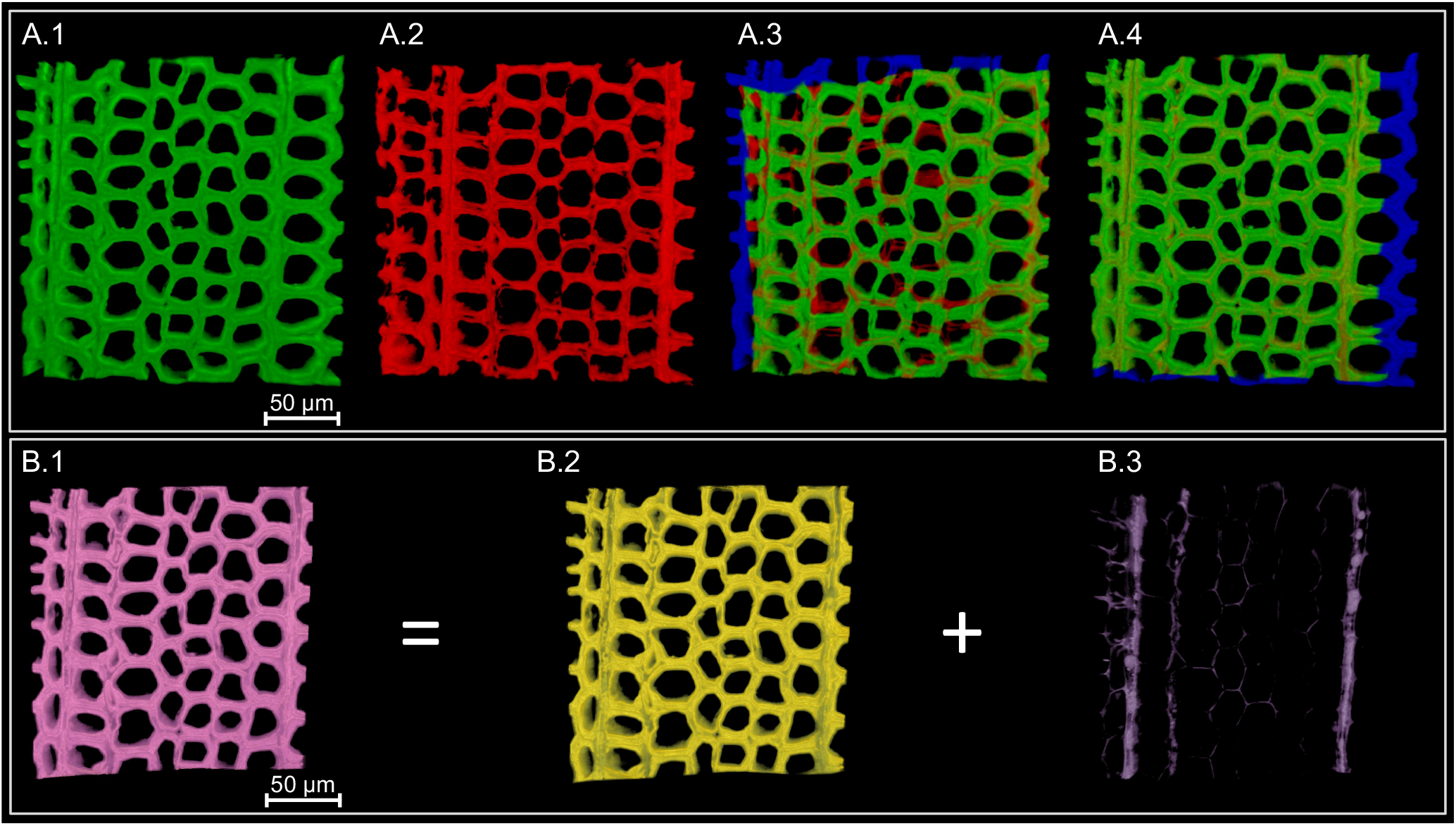
Accurate registration of 3D images of spruce wood under extensive deconstruction conditions providing a voxel resolution spatial deconstruction map. (A.1) Pre-hydrolysis image. (A.2) Hydrolysis phase image after 8 hours of enzymatic deconstruction. (A.3) Unsuccessful direct registration of hydrolysis phase image with pre-hydrolysis image without using divide-and-conquer strategy. (A.4) Successful registration of hydrolysis phase image with pre-hydrolysis image obtained with HydroTrack. (B.1) Consistently imaged region in the frame of pre-hydrolysis image (B.2) Deconstructed voxels after 24 hours of deconstruction of pre-hydrolysis image. (B.3) Remaining voxels of pre-hydrolysis image after 24 hours of enzymatic deconstruction.

HydroTrack’s strategy of dividing time-lapse images into sequential clusters and restricting registrations to images within the clusters, where differences are relatively small and manageable, enables accurate intra-cluster image registration. The successive application of the resulting time-constrained intra-cluster transformations allows to successfully register temporally distance images. Consequently, HydroTrack ensures a higher robustness and reliability in image registration by addressing the inherent limitations of direct registration through a cluster-based indirect registration. The cumulative combining of relatively small, and manageable differences represented through intra-cluster transformations, enables successful image registration even across extensive and unconstrained time intervals. Therefore, HydroTrack allows registration of the hydrolysis phase images, even under extensive deconstruction conditions, with the pre-hydrolysis image (Figure 7.A.4). The subtraction of registered hydrolysis phase image from the prehydrolysis image allows identifying and separating the voxels that have disappeared due to deconstruction from those that remain, even over long time intervals (Figure 7.B). This voxel-by-voxel differentiation reveals the spatial distribution of changes induced by enzymatic deconstruction, mapping out where and to what extent enzymatic hydrolysis has impacted the cell walls. Consequently, a voxel-resolution quantitative 4D perspective (space + time) is offered which presents a remarkably precise and highly detailed mosaic of enzymatic deconstruction of plant cell walls, regardless of the extent of deconstruction.

### 3.3 Adistinct autofluorescence distribution pattern marks cell wall enzymatic hydrolysis

Cell wall tracking enables accurate cell wall autofluorescence intensity dynamics quantification over the course of enzymatic deconstruction. Consequently, autofluorescence intensity dynamics were quantified from collected control and hydrolysis datasets to investigate the effect of enzymatic deconstruction on cell walls. In the control datasets, a first reduction in the autofluorescence intensity values were observed during the initial hours. This was followed by relatively stable peaks and distribution shapes, suggesting minor changes in autofluorescence properties in the absence of enzymatic action (Figure 8.A.1 – A.3). In contrast, the quantification revealed significant shifts in the autofluorescence intensity distributions over the same time intervals for the hydrolysis datasets (Figure 8.B.1 – B.3). Initial changes in distributions were visually comparable to those of the control datasets. However, afterwards, there was a marked decrease in autofluorescence intensity values leading to notable changes in their distributions. This trend was remarkably similar across the different hydrolysis datasets.

**Fig. 8:**
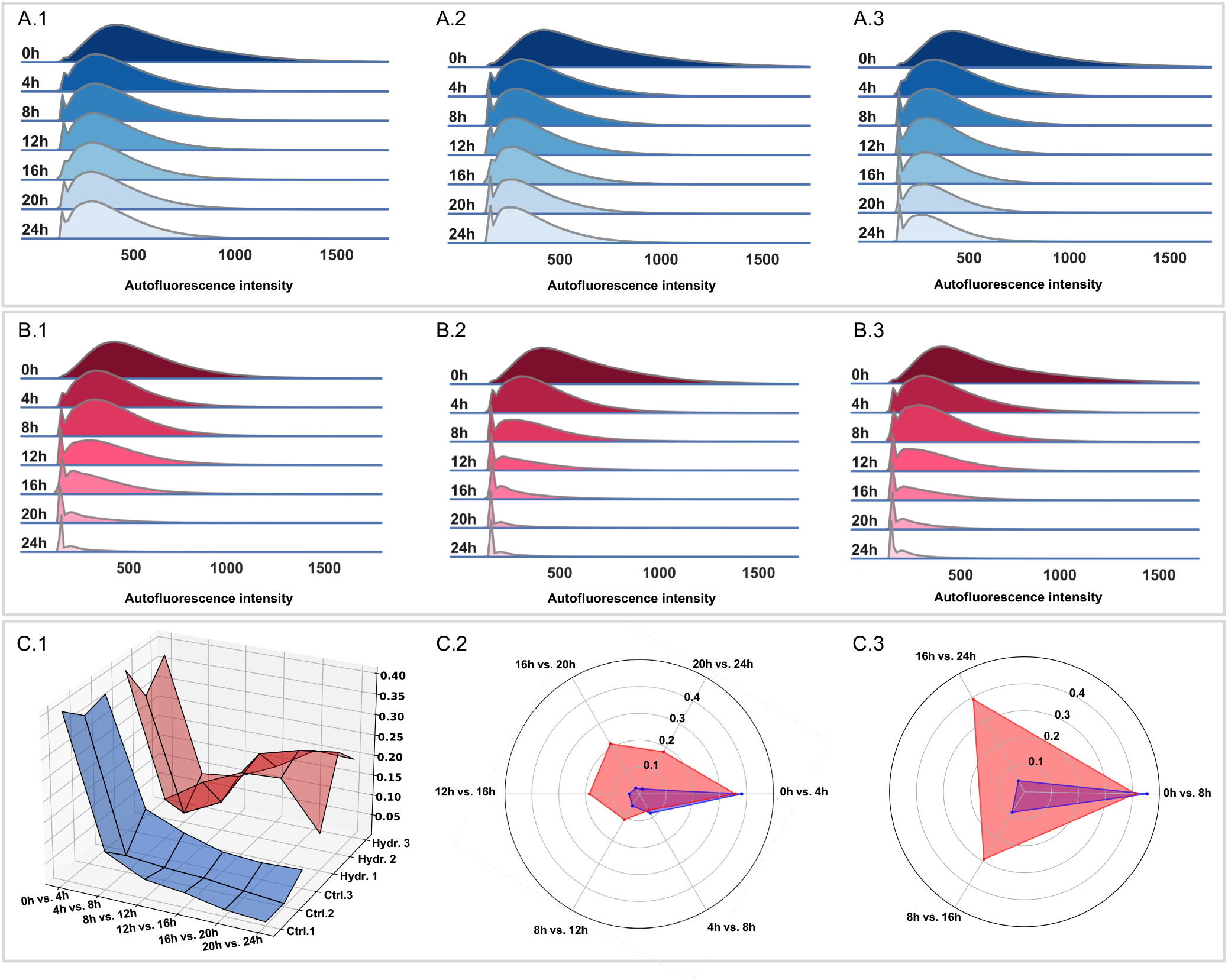
Cell wall autofluorescence intensity dynamics analysis. (A.1 – A.3) Cell wall autofluorescence intensity dynamics illustrated with ridge plots showing the distribution density and its evolution over imaging time, corresponding to the triplicate control time-lapse images, with 4-hour interval. (B.1 – B.3) Cell wall autofluorescence intensity dynamics illustrated with ridge plots, corresponding to the triplicate hydrolysis time-lapse images, with 4-hour interval. (C.1) The Cliff’s delta (*δ*) values for control and hydrolysis datasets represented using 3D surface plots. The height of the surface represents *δ*, with the time intervals and datasets forming the axes. The control datasets are abbreviated as Ctrl. and marked by blue color, and the hydrolysis datasets as Hydr. and marked by red color. (C.2) Average *δ* (effect size) for consecutive distributions with 4-hour interval in control datasets (marked in blue) and hydrolysis datasets (marked in red). (C.3) Average *δ* for consecutive distributions with 8-hour interval in control datasets (marked in blue) and hydrolysis datasets (marked in red). The radar charts highlight the contrast between effect sizes in hydrolysis and control conditions.

To quantitatively analyze temporal evolution of autofluorescence intensity distributions and to assess the magnitude of changes in these distributions, Cliff’s delta, denoted by *δ*, was computed as an effect size [29]. The absolute value of Cliff’s delta indicate the level of effect size, defined as negligible (|*δ*| *<* 0.147), small (0.147 ≤ |*δ*| *<* 0.33), medium (0.33 ≤ |*δ*| *<* 0.474), and large (|*δ*| ≥ 0.474). Effect size is used to quantify the magnitude of difference between two datasets, and provides a measure of the practical significance of the results, beyond the determination of existence of statistical significance. To quantify the cumulative effect of enzymatic hydrolysis on autofluorescence intensity against control conditions, *δ* was first computed between autofluorescence intensity distributions before and after 24 hours of enzymatic hydrolysis (over 24-hour interval). In the hydrolysis datasets large *δ* values were observed with an average of 0.85, indicating significant shifts in autofluorescence intensity distributions. In contrast, the control datasets showed lower *δ* values with an average of 0.56. The differences between autofluorescence intensity distributions over the 24-hour interval, for both control and hydrolysis datasets, were statistically significant (Kruskal-Wallis test, *p*-value *<* 10^*−*5^).

To quantitatively examine the incremental effects of enzymatic hydrolysis on cell wall autofluorescence and investigate the intermediate stages of autofluorescence intensity changes, the time interval was then reduced to a 4-hour interval. The differences between successive autofluorescence intensity distributions in control and hydrolysis datasets, were statistically significant (Kruskal-Wallis test, *p*-value *<* 10^*−*5^). *δ* was then computed between pairs of successive autofluorescence intensity distributions (Figure 8.C.1). For control datasets, the *δ* values were medium during the first time interval (0h - 4h), with an average of 0.38 (specifically, 0.41, 0.36, 0.38, for the three control datasets). Over the subsequent intervals (4h – 8h, 8h - 12h, …, 20h - 24h), *δ* values were negligible with an average below 0.1, indicating minor changes in autofluorescence distributions in the absence of enzymatic hydrolysis. For hydrolysis datasets, *δ* values were similarly medium in the first interval (0h - 4h) with an average of 0.36 (specifically, 0.4, 0.3, 0.37, for the three hydrolysis datasets) and also negligible over (4h - 8h). Over the next time intervals (8h - 12h, …, 20h - 24h), average *δ* ranged from 0.11 to 0.22, which were significantly higher than those computed for the control datasets during the same time intervals (Figure 8.C.2). The similarity of *δ* values between control and hydrolysis datasets during the initial 8 hours led to using an 8-hour time-interval resolution. With 8-hour time interval, the differences between pairs of successive autofluorescence intensity distributions in control and hydrolysis datasets, were statistically significant (Kruskal-Wallis test, *p*-value *<* 10^*−*5^). After the initial 8 hours, hydrolysis datasets showed higher *δ* values compared to control datasets (exhibiting negligible *δ* values) highlighting the enzymatic impact (Figure 8.C.3). To evaluate the cumulative impact of enzymatic hydrolysis against control conditions during post-initial time intervals, *δ* was computed between autofluorescence intensity distributions at 4h and 24h and also between distribution at 8h and 24h. Over the 8h – 24h interval, the average *δ* was 0.65 and 0.14 for the hydrolysis datasets and control datasets respectively. Similarly, over the 4h – 24h interval, the average *δ* was 0.7 and 0.22 for the hydrolysis datasets and control datasets respectively.

The quantification of magnitude of difference between pairs of distributions with different temporal resolutions reflecting both incremental and cumulative effects of enzymatic hydrolysis, confirmed the initial visual observations of contrasting patterns of autofluorescence intensity distributions between hydrolysis and control datasets. This contrasts is further emphasized in the radar charts shown in (Figure 8.C.2 & C.3), which illustrate the magnitude of autofluorescence intensity changes between hydrolysis and control datasets. Together, these results demonstrated that the hydrolysis datasets exhibit a distinguishing pattern of autofluorescence intensity distribution shifts which effectively differentiating them from the autofluorescence intensity distributions of the control datasets, demonstrating that autofluorescence intensity is an distinctive indicator characterizing the cell wall enzymatic deconstruction.

### 3.4 Accessible surface area reduction lags behind volume decrease in cell walls, revealing asynchronous effects of deconstruction

The HydroTrack pipeline identifies individual cells from time-lapse images of cell wall deconstruction, which enables investigation of the impact of enzymes on cell wall structural and morphological parameters. The application of HydroTrack to collected datasets generated segmented time-lapse image datasets with total number of 6505 and 6699 segmented cells in control and hydrolysis datasets respectively. Consequently, the temporal evolutions of two critical structural parameters, cell wall accessible surface area and cell wall volume, for both control and hydrolysis datasets were quantified (Figure 9.A & B). The quantifications were limited to consistently imaged region across time-lapse images to ensure comparability of the quantifications over time. Therefore, cell walls that were captured at some time, but partially or completely absent at others, were excluded from the analysis. The cell wall accessible surface area can be considered as a cell scale analogue to the accessible surface area of the polysaccharide matrix within the cell wall. This parameter is crucial, as a higher accessible cell wall surface area is associated with increased accessibility to polysaccharides, therefore can significantly influence the yield of enzymatic deconstruction of plant cell wall [30, 38]. The cell wall volume is also a key parameter which provides a cell scale indicator of the enzymatic deconstruction efficiency [38].

**Fig. 9:**
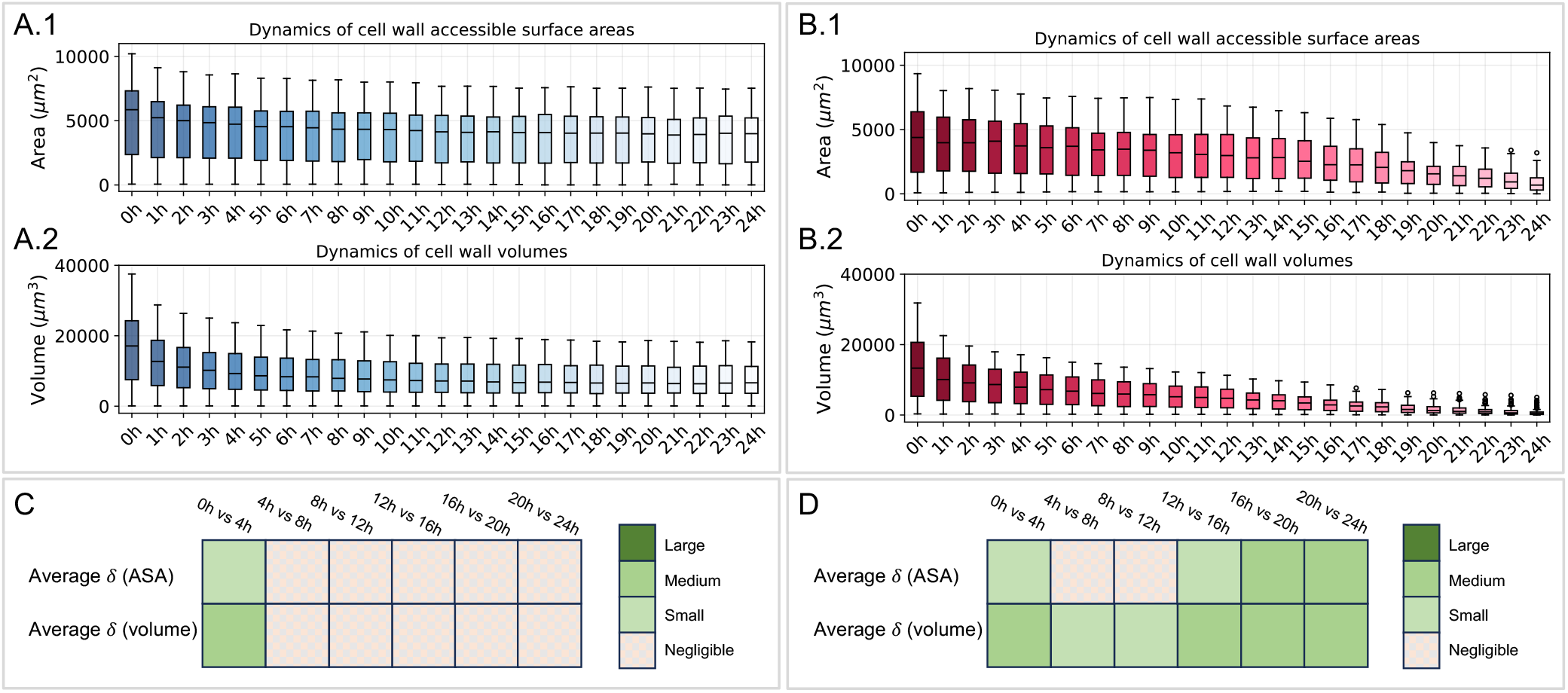
Cell wall accessible surface area and cell wall volume dynamics analysis. (A.1 & A.2) Dynamics of accessible surface area and cell wall volume, respectively, in a representative control dataset, with 1-hour interval. (B.1 & B.2) Dynamics of accessible surface area and cell wall volume, respectively, in a representative hydrolysis dataset, with 1-hour interval. (C) Average cliff’s *δ* (effect size) for consecutive cell wall accessible surface area (ASA) and cell wall volume distributions with 4-hour interval for the control datasets. (D) Average cliff’s *δ* for consecutive cell wall accessible surface area (ASA) and cell wall volume distributions with 4-hour interval for the hydrolysis datasets. In the boxplots, the median values are indicated by the black lines and circles indicate the outliers. The colors bars indicate *δ* values defined as negligible (*δ <* 0.147), small (0.147 ≤ *δ <* 0.33), medium (0.33 ≤ *δ <* 0.474), and large (*δ* ≥ 0.474). For the control datasets, the number of cells in the consistently imaged region within the first acquisitions were 75 (displayed control dataset), 91, and 96. The total number of cells in the consistently imaged regions across all time points were 1875 (displayed control dataset), 2225, and 2375 for the triplicate control time-lapse images. For the triplicate hydrolysis time-lapse images, the number of cells in the prehydrolysis consistently imaged region were 96 (displayed hydrolysis dataset), 95, and 90 with the total number of cells in the consistently imaged regions across all time points being 2324 (displayed hydrolysis dataset), 1986, and 2222.

After quantifying cell wall volumes and accessible surface areas, their distributions were analyzed at 4-hour intervals to evaluate the magnitude of differences (using Cliff’s delta) (Figure 9.C & D). Control datasets exhibited non-negligible average *δ* values between distributions only over the initial time interval (0h - 4h) for both cell wall volume and accessible surface area. Over the later time intervals, average *δ* values became negligible, indicating that cell wall volume and surface area remain relatively stable after initial hours. Consistent with this, statistically significant differences between distributions were only conclusively observed over 0h - 4h interval (Kruskal-Wallis test, *p*-value *<* 0.05).

The hydrolysis datasets exhibited non-negligible average *δ* values between distributions over the initial 4 hours in cell wall volume and accessible surface area, similar to control datasets. However, over later time intervals, non-negligible average *δ* values between cell wall volume distributions were consistently observed. In contrast, two distinct trends were observed for cell wall accessible surface area during intermediate (including 4h - 8h and 8h - 12h intervals) and final (including 12h - 16h, …, 20h - 24h intervals) intervals: negligible *δ* values over intermediate intervals and significant distribution shifts over the final intervals, characterized by small to medium effect sizes. In line with this, statistically significant differences between accessible surface area distributions were only conclusively observed over final intervals (Kruskal-Wallis test, *p*-value *<* 0.05).

These results indicated that after the initial distribution shifts, the volume of cell walls were more consistently impacted in the early stages of the enzymatic deconstruction compared to cell wall accessible surface area. For cell wall accessible surface area, significant shifts are not as immediately observable as they are for cell wall volume but become more noticeable in later intervals. Together, theses results revealed a different temporal evolution of cell wall volume and cell wall accessible surface area, demonstrating an asynchronous impact of hydrolysis on cell wall structural parameters with leading volume reduction and following accessible surface decrease.

### 3.5 New perspectives to assess enzymatic deconstruction dynamics

This study provides a robust automatic pipeline, named HydroTrack, which addresses a comprehensive set of challenges encountered in quantitative microscale study of cell wall enzymatic deconstruction using time-lapse imaging. By successfully handling complexities due to sample displacement, autofluorescence intensity reduction and cell wall deformation occurring during imaging of cell wall enzymatic deconstruction, HydroTrack tracks cell walls deconstruction and enables quantification of autofluorescence intensity dynamics. The cell wall resolution segmentations generated by HydroTrack allows to investigate the cell scale impact of enzymatic hydrolysis. To the best of our knowledge, HydroTrack is the only automated pipeline which allows the study of cell wall modifications at cell and tissue scales under high deconstruction conditions. As shown before, accurate tracking of cell wall autofluorescence intensity dynamics and cell wall morphometrics enables assessment of enzymatic deconstruction kinetics and efficiency of enzymatic hydrolysis and can serve as an indicator of enzymatic deconstruction [38]. This makes HydroTrack an essential pipeline in the precise quantitative analysis of cell wall deconstruction processes. Importantly, the quantifications revealed distinct patterns of autofluorescence intensity shifts during cell wall enzymatic deconstruction, effectively differentiating them from the autofluorescence intensity distributions of the control datasets. These results underscore the potential of autofluorescence intensity distributions to serve as a distinguishing indicator of cell wall enzymatic deconstruction. Quantification of temporal dynamics of cell wall morphometrics provided critical insights into the sequence of structural changes within the cell wall and revealed asynchronous dynamics of cell wall volume and cell wall accessible surface area. Taken together, this study offers a robust approach to observe and analyze the cell wall deconstruction with novel findings, enabling the exploration of previously unexamined aspects of cell wall deconstruction in unprecedented detail. By comprehensively addressing the challenges of quantitative microscale investigation of cell wall enzymatic hydrolysis, the pipeline ensures a broad applicability to diverse biomass species and pretreatment methods. This approach not only enhances the fundamental understanding of deconstruction but also provides methodological advancements with potential significant implications by enabling robust cell scale assessment of enzymatic deconstruction dynamics which can be used to evaluate and optimize deconstruction efficiency and tailored enzyme cocktails and pretreatment strategies. Thus, our study provides a pathway to bridge fundamental research with industrial application, enabling more efficient conversion of plant cell wall into bioproducts.

## 4 Conclusions

A sustainable transformation of plant cell wall requires a deeper, integrated understanding of enzymatic deconstruction. This study presents a novel and robust 4D quantitative pipeline specifically designed to investigate plant cell wall enzymatic hydrolysis under high deconstruction conditions. Application of the pipeline to pretreated spruce wood revealed a distinct autofluorescence distribution pattern of cell wall hydrolysis and an asynchronous impact of hydrolysis on cell wall morphometrics. This study provides novel insights into enzymatic deconstruction of cell walls and novel methodological advances paving the way for a more efficient conversion of plant cell walls into bioproducts.

## CRediT authorship contribution statement

Solmaz Hossein Khani: Conceptualization, Data curation, Formal analysis, Investigation, Methodology, Software, Validation, Visualization, Writing – original draft, Writing – review and editing. Khadidja Ould Amer: Data curation, Methodology, Writing – original draft. Noah Remy: Methodology, Investigation, Writing – original draft. Berangère Lebas: Methodology, Investigation, Writing – original draft. Anouck Habrant: Methodology, Investigation, Writing – original draft. Ali Faraj: Methodology, Software, Writing – review and editing. Grégoire Malandain: Investigation, Methodology, Software, Writing – review and editing. Gabriel Paës: Conceptualization, Formal analysis, Funding acquisition, Investigation, Methodology, Project administration, Resources, Supervision, Validation, Writing – original draft, Writing – review and editing. Yassin Refahi: Conceptualization, Data curation, Formal analysis, Funding acquisition, Investigation, Methodology, Project administration, Resources, Software, Supervision, Validation, Visualization, Writing – original draft, Writing – review and editing

## Acknowledgments

We thank Antoine Margeot and Simon Arragain from IFPEN (Rueil-Malmaison, France) for providing the enzymatic cocktail. We thank Juliette Floret for determining enzymatic activity and François Gaudard for his assistance with the preparation and characterization of spruce tree wood samples.

